# Capacity for compensatory cyclin D2 response confers trametinib resistance in canine mucosal melanoma

**DOI:** 10.1101/2025.04.24.650512

**Authors:** Bih-Rong Wei, Vincenzo Verdi, Shuling Zhang, Beverly A. Mock, Heather R. Shive, R. Mark Simpson

## Abstract

**Background/objective:** Mucosal melanoma (MM) is a poorly responsive, rare and aggressive subtype with paucity of targetable recurrent driver mutations, although Ras/MAPK and PI3K/AKT/mTOR signaling pathway activations are common. Eventual tumor evasion of targeted therapy continues to limit treatment success. Adequate models are necessary to address therapeutic resistance. The relatively greater incidence of naturally occurring MM in dogs, as well as its comparable clinical and pathological characteristics to human MM, represents a promising opportunity for predictive patient modeling. Resistance-promoting crosstalk between Ras/MAPK and PI3K/AKT/mTOR signaling under trametinib inhibition of MEK was studied in a canine MM model. Emphasis was placed on the suppressive effect of trametinib on cell cycle entry and its potential role in drug resistance.

**Methods:** D-type cyclins were investigated using five MM cell lines exhibiting differential sensitivities to trametinib. Drug-treated cells were analyzed for signaling pathway activation, proliferation, survival, cell death, and cell cycle in the context of D-type cyclin expression. Cyclin D2 expression was manipulated using siRNA knock down or inducible recombinant overexpression.

**Results:** With diminished cyclin D1 under trametinib treatment, relatively trametinib-resistant MM cells exhibited capacity to upregulate cyclin D2, which promoted proliferation, whereas sensitive cells did not similarly respond. Inhibition of the compensatory cyclin D2 response restored sensitivity to resistant cells. Induced cyclin D2 overexpression promoted survival to otherwise trametinib-sensitive MM cells that did not exhibit capacity to upregulate endogenous cyclin D2. PI3K/AKT/mTOR signaling upregulation under trametinib was suppressed by mTORC1/2 inhibition, which similarly diminished cyclin D2 response.

**Conclusion:** The compensatory switch from preferential reliance on cyclin D1 to D2 appears to play a role in MM resistance to MEK inhibition.

## Introduction

Mucosal melanoma (MM) is a rare and aggressive melanoma subtype that arises in the mucous membranes lining various internal body cavities such as the nasal passages, oral cavity, gastrointestinal tract, and genitourinary tract. Distinct from cutaneous melanoma, MM accounts for only about 1% of all melanomas and is not associated with cutaneous exposure to ultraviolet (UV) light as a risk factor (1). It is often diagnosed at advanced stages due to anatomical inaccessibility and nonspecific early symptoms, characteristics contributing to its poor prognosis. The five-year survival rate for MM remains below 20%, compared to 90% for cutaneous melanoma (2, 3). This underscores the need for improved diagnostic and therapeutic strategies. The rare incidence and limited sources of human MM cell lines and tissues for research have hampered progress developing treatments. Canine MM shares conspicuous parallels with human MM (4–6), and investigations of naturally occurring MM in pet dogs can serve as an ideal surrogate comparative and translational model for human MM (5, 7). Clinically, canine MM mirrors human MM’s invasive growth, frequent metastasis (e.g., to lymph nodes and lungs), poor therapeutic response and high recurrence rate despite surgery, radiation therapy or both (8). Expediently, the spontaneous occurrence of canine MM, in an immunocompetent host, with its relative greater incidence in dogs than in humans (up to 100,000 canine diagnoses annually), offers a naturally occurring patient model to develop and refine therapies for both canine and human MM, bridging preclinical and clinical gaps (9–11).

In contrast to human cutaneous melanoma, there is a lack of recurrent dominant driver mutations to guide selection of prevailing targeted therapy for both human and canine MM (12–14). In human MM, 20-40% cases harbor c-KIT mutations (12, 15), however, only approximately 15-21% of these cases include activating mutations in c-KIT (L576P, K642E) (15). The therapeutic response to c-Kit inhibitors is varied, and less than ideal (16). Similarly, c-Kit mutations are rare in canine MM (<10%) (6, 17) and c-kit inhibitors have shown limited efficacy in canine experimental models (18). Additionally, BRAF mutation is rare (8%) and NRAS mutations (10-25%) vary in frequency (12, 19, 20). Generally, human and canine MM both feature a low overall mutational burden (human: 2-5 m/Mb; canine: 1-4 m/Mb), in contrast to human cutaneous melanoma (10-20 mutations/Mb) (13, 21). Nevertheless, targeting signaling pathways activated in canine and human MM, such as Ras/MAPK and PI3K/AKT/mTOR, remains a viable option due to their frequent activation and vital role in supporting tumor growth (4, 22). Targeting Ras/MAPK and/or PI3K/AKT in canine MM has been shown to inhibit specific target signal transduction while inducing cell death, cell cycle arrest, and suppressing tumor growth and metastasis in mouse xenograft models (23).

The Ras/ERK pathway is a pivotal mitogenic cascade that drives cell proliferation by modulating cyclin D expression, a critical regulator of the G_1_-to-S phase cell cycle transition via CDK4/6 activation (24). The extracellular signal-regulated kinase (ERK), a key component of Ras signaling, is sequentially activated by Ras and mitogen-activated protein kinase kinase (MEK), which leads to enhanced cyclin D1 (CCND1) transcription by phosphorylating transcription factors such as c-Myc and AP-1, thereby transducing extracellular signals into cell cycle progression (25). Cyclin D1 expression, augmented in several human cancer types has been found in over 60% of human and canine MM (26, 27). Inhibition of MEK through allosteric binding by small molecule inhibitors such as trametinib, prevents MEK phosphorylation and subsequent activation of ERK, leading to suppression of cyclin D1 level (28–30).

Trametinib has demonstrated an objective response rate of approximately 22% in BRAF-mutated human cutaneous melanomas as a monotherapy, with a median progression-free survival (PFS) of 4.8 months (31). Despite trametinib’s initial efficacy, most patients eventually develop resistance (32, 33), which underscores the need for research into resistance mechanisms and countermeasures. The mechanisms of resistance are diverse and complex, including target-specific alterations like second-site mutations, amplification of the target kinase, and activation of alternative signaling pathways (34). Additionally, some 10% of melanoma patients with Ras/MAPK activation fail to respond to targeted inhibition (BRAF and MEK) and are considered intrinsically resistant (35).

Counteracting oncology drug resistance requires a deeper understanding of these mechanisms in the context of disease. Such insights will be instrumental in designing better drug regimens to improve patient outcomes. Investigating mechanisms of drug resistance in canine MM models, which exhibit high fidelity for human MM, can inform means to circumvent resistance to targeted therapies, both for their intrinsic value for dogs with spontaneous MM and for comparative translational potential benefit in human MM (36). Despite trametinib’s faculty for reducing cyclin D1 level upon treatment, we noted some canine MM cells were able to proceed with cell cycle progression and proliferate under trametinib treatment in this study. Considering the essential function of D-type cyclins in initiating cell cycle, we investigated the role of other cyclin D family members, i.e., cyclin D2 and cyclin D3, in canine MM resistance to trametinib.

## Materials and methods

### Cell lines, cultures, and cell fate assays

Canine mucosal melanoma cell lines were originally derived ex vivo from canine mucosal melanomas patients and provided by Dr. Michael Kent at University of California, Davis (UCDK9M1 (M1), UCDK9M2 (M2), UCDK9M3 (M3), UCDK9M4 (M4), UCDK9M5 (M5)), and Drs. Jared Fowles and Dan Gustafson of Colorado State University, Fort Collins (Jones) (37, 38). Cells were cultured in DMEM with 10% FBS in 5% CO_2_ at 37°C. All cell lines were routinely tested for mycoplasma contamination.

M5 cells overexpressing cyclin D2 (M5D2) were generated by co-transfecting M5 cells with XLone-Puro CCND2-T2A-luciferase-P2A-eGFP, a gift from Xiaoping Bao Lab, Purdue University (Addgene plasmid # 179843) and Super PiggyBac transposase expression vector (System Biosciences, PB210PA-1) at 1:2.5 ratio using jetPRIME transfection reagent (PolyPlus.Poluplus-transfection.com). The transfected cells were cultured under puromycin (2.5 ug/ml) selection for two weeks. To induce cyclin D2 expression, 0.1 µg/ml doxycycline (Sigma-Aldrich, Inc. St. Louis, MO 63178) was added to the culture medium one day before experiments.

The trametinib resistant M1 (M1T^R^) and M2 (M2T^R^) cells were generated by culturing cells in increasing concentration of trametinib from 16 nM to 10 µM in a period of 2 – 4 weeks. The M1T^R^ and M2T^R^ cells were then maintained in culture medium containing 2 µM trametinib. Cell titers were determined by MTS assay using CellTiter 96® AQueous Non-Radioactive Cell Proliferation Assay (MTS, Promega) following manufacturer’s protocol. A total of 3×10^3^ cells were seeded in each well of 96 well plates. At the end of treatment, the MTS solution was added to the cells at 1:5 ratio and incubated for 1.5 hours. A microplate reader (SPARK, TECAN) was used to measure the OD at 490 nm.

Cell death was evaluated using Incucyte® Cytotox Dyes and Incucyte® Annexin V Dyes (Sartorius). A total of 0.5 - 1×10^4^ cells were seeded in each well of 24 well plates 24 hours prior to incubation with the drugs and the dyes for the duration of the experiments. Cytotox Dye was used at a final concentration at 250 nM. Annexin V dye was diluted at 1:200 in culture medium. The cell images were acquired every 2-3 hours using Incucyte S3 with 10x objectives (Sartorius). Image analyses, confluency, cytotoxicity and Annexin V labeling were quantified using Incucyte software.

### Western blotting

Cell lysates of cultured cell lines were generated by lysing cells in cell lysis buffer (Cell Signaling) for 20 minutes on ice. Cell lysates were cleared by centrifugation at 18,000 X g for 10 minutes. The cleared cell lysates were separated on 4-20% Tris-glycine gels (Invitrogen) and transferred onto PVDF membranes (Bio-Rad). Membranes were probed with primary antibodies (Supplemental Table S1) followed by respective horseradish peroxidase (HRP)-conjugated secondary antibodies (Jackson Immunoresearch). Immunoreactive bands were detected using chemiluminescence and GE Amersham Imager 600 (Amersham/Cytiva). The intensities of the signals were quantified using ImageJ.

### Cell cycle flow cytometry

To analyze cell cycle, cells were plated one day before treatment application. 1×10^6^ cells were then washed with PBS at designated time and fixed with 70% ethanol. RNA was removed by adding 100 units of RNase A (Sigma) and DNA was stained with 50 ug/ml propidium iodide. Cell cycle was analyzed using LSRFortessa (BD Biosciences) and ModFit LT 5.0 software.

### siRNA knock down

For siRNA knockdown (KD) experiments, cells were plated one day prior to transfection and transfected with 30 nM siRNA targeting CCND1 or CCND2 or control siRNA (siGENOME Non-Targeting Control siRNAs, 6505, Dharmacon) using DharmaFECT 1 transfection reagents (Dharmacon). After 24-hour incubation, the siRNA was removed and fresh medium containing trametinib was added to cells for 24 – 48 hours. Multiple siRNA constructs were tested for each gene (Supplemental Table S1) and showed comparable efficiency (Supplemental Figure S1). A mixture of equal quantities of siRNAs were used in the subsequent experiments.

### RNA extraction and quantitative RT-PCR analyses

Total RNA was isolated using RNeasy Plus Kits (Qiagen) following manufacturer’s manual. RNA quantity and quality were assessed by spectrophotometry (NanoDrop ONE,122 Waltham, MA, USA). Total RNA was reverse-transcribed using High Capacity cDNA Reverse Transcription Kit (Applied Biosystems, Cat.# 4368814;) according to the manufacturer’s protocol. Quantitative real time PCR (qRT-PCR) analysis was performed using AzuraView™Green Fast qPCR Blue Mix (Cat.# AZ-2350; Azura Genomics) and BIO-RAD CFX connect real time system (Bio-rad.com). Gene expression was normalized to the expression of the housekeeping gene, b-actin. Primer sequences are listed in Supplemental Table S1.

### EdU incoporation

EdU incorporation was performed following the siRNA and/or drug treatment using Click-iT imaging kit (Invitrogen). Briefly, cells were pulsed with 10 uM EdU for 1.5 hours prior to the fixation followed by labeling of incorporated EdU according to the manufacturer’s protocol. Cyclin D1 or cyclin D2 immunofluorescent staining was performed subsequently. Briefly, following EdU labeling, cells were incubated with primary antibodies against cyclin D1 or cyclin D2, Alexa-594 conjugated goat anti-rabbit IgG secondary antibody (1:200 dilution) (ThermoFisher) and DAPI (1:1000 dilution) (ThermoFisher) for nuclear counterstaining. The images were captured using Zeiss Imager M2 and Zeiss Zen software (Zeiss). The % EdU positive cells were calculated by number of EdU positive cells/total (DAPI) cells in each image. Images of ten to twenty-five random fields were taken for each treatment group.

### Colony formation assay

Cells were seeded in 24-well plates at a density of 2,000 cells per well and cultured with trametinib for 1 week. Following the incubation period, the cells were fixed with 20% methanol, and 0.5% crystal violet solution was used to stain and visualize the colonies. Images were captured using Leica M165 FC stereo microscope. Control cells were treated with an equivalent volume of DMSO.

### Reagents

Antibodies against p-AKTSer473, p-S6Ser235/236, p-ERK 1/2, cyclin D2 were obtained from Cell Signaling Technologies (Danvers, MA). Anti-cyclin D1 and cyclin D2 were from Abcam (Cambridge, MA). Anti-b-actin was obtained from Sigma-Aldrich (St. Louis, MO). Detailed information is listed in Supplemental Table S1.

Sapanisertib, colbimetinib, selumetinib, and binimetinib were purchased from MedChem Express (Monmouth Junction, NJ). Trametinib was purchased from ChemieTek (Indianapolis, IN). Both drugs were dissolved in DMSO to make a stock solution of 10 μM. When treating cells, the drugs were diluted in DMSO to 1000x designated concentration and added to cell culture medium at 1:1000 (v:v) dilution.

### Statistical Analysis

Statistical analyses were conducted using GraphPad Prism version 10.3.1 or Microsoft Excel. A two-sample unequal variance t-test was performed to compare two groups. P values < 0.05 were considered statistically significant.

## Results

### Differential intrinsic resistance to trametinib in a series of mucosal melanoma cell lines

Five canine mucosal melanoma (MM) cell lines were treated with trametinib over a range of concentrations, and cell titers were determined by MTS assays after 72 hours. The cell titers at the end of 72 hour treatment revealed differing levels of trametinib sensitivity (Figure 1A and Supplemental Table S2). M1 and M2 MM cell lines were relatively resistant to trametinib, while M5 and Jones were more sensitive to trametinib at the concentrations tested. The M3 cell line was intermediate in sensitivity. This relative sensitivity among the cell lines was sustained across a range of trametinib concentrations (Figure 1A). The differential sensitivity of MM cells to trametinib was further reflected in a colony formation assay (Figure 1B). We also assessed cell cycle in both relatively resistant (M1) and sensitive (M5) MM cells at various trametinib concentrations (0.1, 0.5 and 1.0 µM). In contrast to control cultures lacking trametinib, fractions of cells in G_1_ cell cycle phase were significantly increased and G_2_ and S phase cell populations were decreased when exposed to trametinib (Figure 1C). An EdU incorporation assay indicated that most cell lines retained the ability to proliferate, although at lower rate compared to untreated control cells (Figure 1D). In addition, trametinib treated M1 and M5 cells underwent cell death to degrees mirroring their relative inherent sensitivities (Figure 1E).

**Figure 1.**
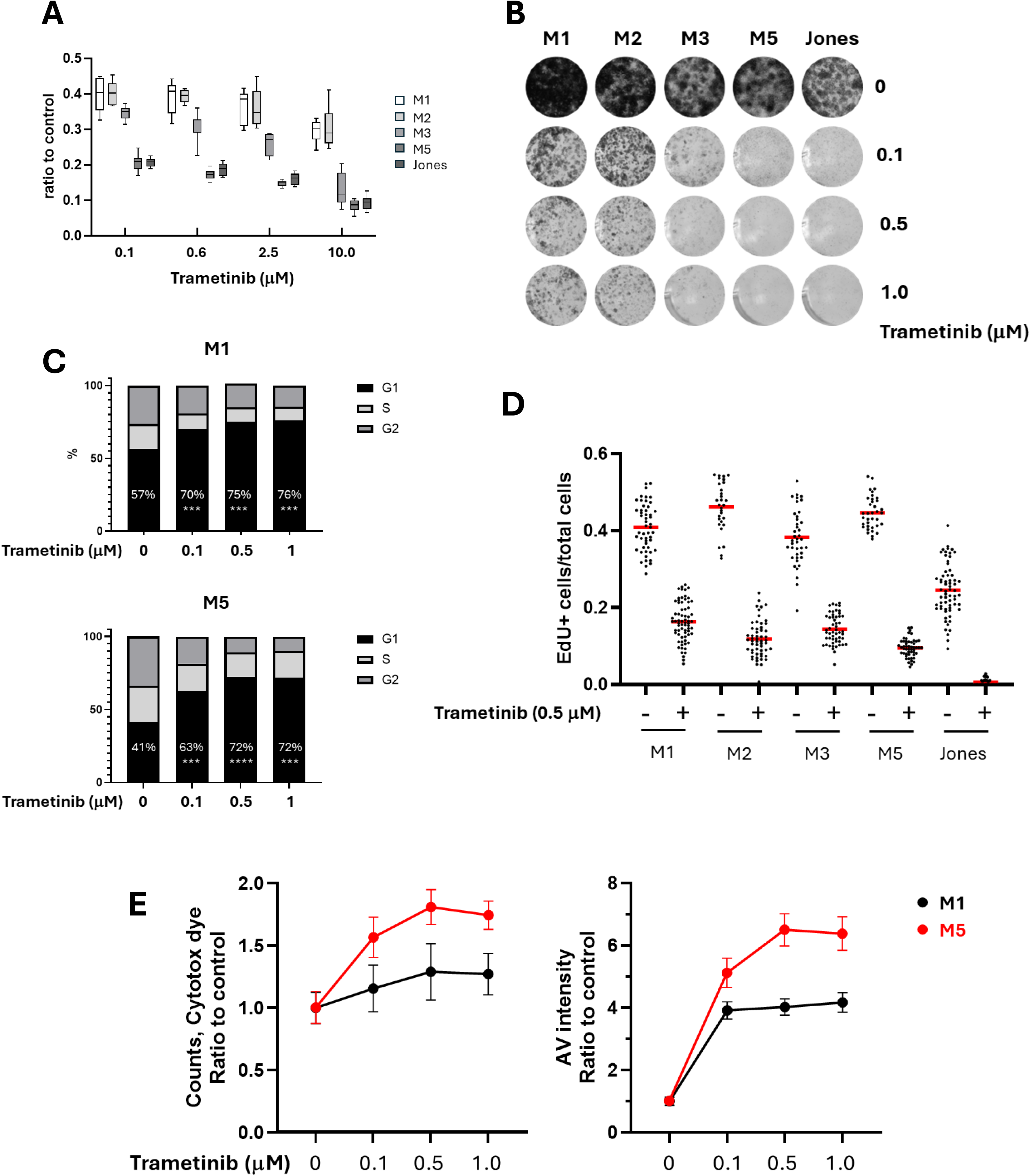
A series of canine mucosal melanoma cell lines exhibited differing sensitivities to acute trametinib exposure. A) Cells were treated with serial concentrations of trametinib for 72 hours and cell viability titers were determined by MTS assay. B) Colony formation assays of canine MM cells treated with trametinib indicate outcomes that parallel viability relationships among cell line series. C) Trametinib induced G1 cell cycle arrest in both trametinib resistant (M1) and sensitive (M5) cells. Cell cycle analyses were performed on M1 and M5 cells treated with trametinib for 24 hours. D) Cells surviving trametinib treatment retain proliferative capacity. Ratios of cells expressing EdU incorporation to total cells for five MM cell lines, both with and without the addition of trametinib. E) Trametinib sensitive M5 cells (red line) exhibited higher rate of cell death compared to trametinib resistant M1 cells (black line). M1 and M5 cell were treated with trametinib, and the incidence of cell death was evaluated up to 72 hours using Incucyte cytotox Dye and Annexin V labeling assays. The labeling at 72 hours was illustrated to represent the degree of cell death at different trametinib concentrations, referenced to nontreated control conditions.

### Cyclin D expression profiles in canine MM cells

Cyclin D plays a pivotal role in cell proliferation by driving progression through the G_1_ phase and enabling the G_1_ to S cell cycle transition. With evidence of differing cell fate under trametinib treatment (Figure 1) and considering that subfamily members of cyclin D have varying roles in driving cell cycle progression, we investigated whether the expression profile of cyclin Ds in canine MM cells was associated with their response to trametinib. All MM cell lines expressed cyclin D1 and cyclin D2 at both transcriptional and protein levels (Figure 2A, B). In contrast, minimal cyclin D3 expression was detected by quantitative PCR (Figure 2A), and cyclin D3 protein was not evident. Nuclear labeling of cyclin D1 and cyclin D2 suggested cells utilize both cyclin Ds during the G_1_ phase (Figure 2C). The non-overlapping pattern of EdU positive cells (S phase) and nuclear cyclin D positive cells (G_1_ phase) corresponds to the role of cyclin D in cell cycle. Endogenous levels of cyclin D expression, therefore, did not appear to parallel intrinsic trametinib sensitivity.

**Figure 2.**
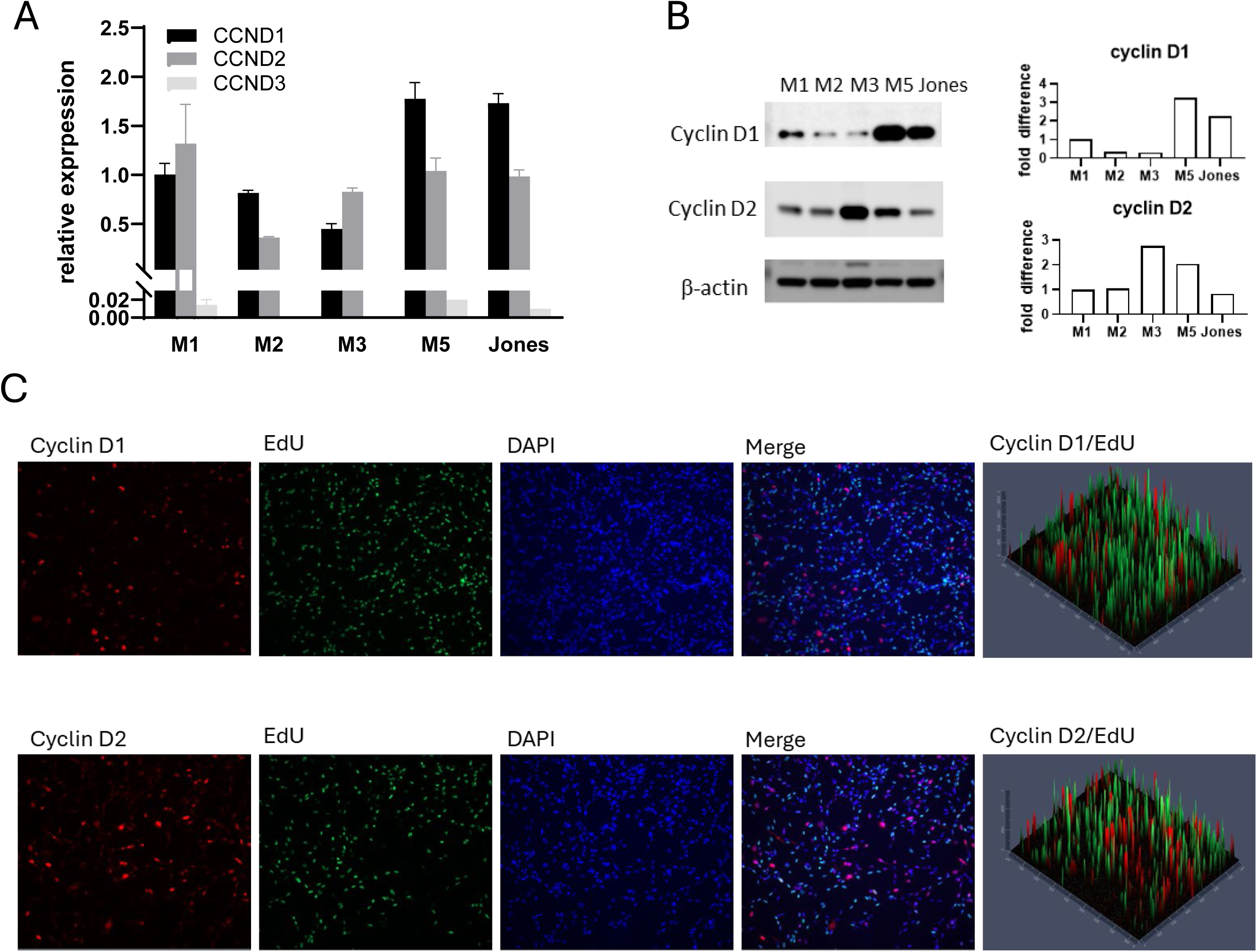
Canine MM cell lines expressed variable levels of cyclin Ds, primarily both cyclin D1 and D2. A) Quantitative PCR detection of cyclins D1, D2 and D3 mRNA levels. B) Cyclins D1 and D2 protein levels were assessed in canine MM cells in linear growth phase. C) Immunofluorescent labeling of cyclin D1 and cyclin D2 in M5 cells. All MM examined expressed both cyclin D1 and cyclin D2. Representative immunofluorescent labeling of M5 cells are shown.

### Trametinib-resistance projected by compensatory cyclin D2 response

Trametinib reduces the expression of cyclin D1 in canine melanoma cells (29), as well as cell lines of human non-small cell lung cancer (39), melanoma (40, 41), and colon cancer (42). Whether trametinib influences the expression of cyclin D2 and cyclin D3 is not well studied. Certain cell types are known to exhibit preferential expression of a particular D-type cyclin, and in the absence of a dominant subtype another D-type cyclin may compensate (43). We next examined how trametinib affected the expression of cyclin Ds in the series of canine melanoma cells. Cells were treated with 0.1 or 1 µM trametinib for 24 and 48 hours. The expression of cyclin Ds was documented by quantitative PCR for transcription and by western blots for protein levels (Figure 3A, B). Upon trametinib treatment, cyclin D1 was diminished in all five MM cell lines. The cyclin D2 responses, however, were noteworthy for their corresponding differences among cells across the range of trametinib sensitivities. The cyclin D2 level in the trametinib-sensitive cell lines, M5 and Jones, was decreased along with cyclin D1. By contrast, relatively resistant MM cells (M1 and M2) up-regulated cyclin D2 expression as cyclin D1 was downregulated. In M3 cells, the intermediate responder to trametinib, a delayed and more muted cyclin D2 response was observed; cyclin D2 remained comparable with or without trametinib treatment at 24 hours, while at 48 hours cyclin D2 level was slightly increased (Figure 3A). Cyclin D3 levels were relatively low compared to cyclin D1 and cyclin D2, with or without trametinib treatment, in all five MM cells (Supplemental Figure S2). The reciprocal expression of cyclin D2 upon cyclin D1 reduction during MEK inhibition in MM did not appear to be trametinib specific. The effects of other MEK inhibitors, cobimetinib, binimetinib, and selumetinib, were similar to trametinib on M1 MM cells (Figure 3C).

**Figure 3.**
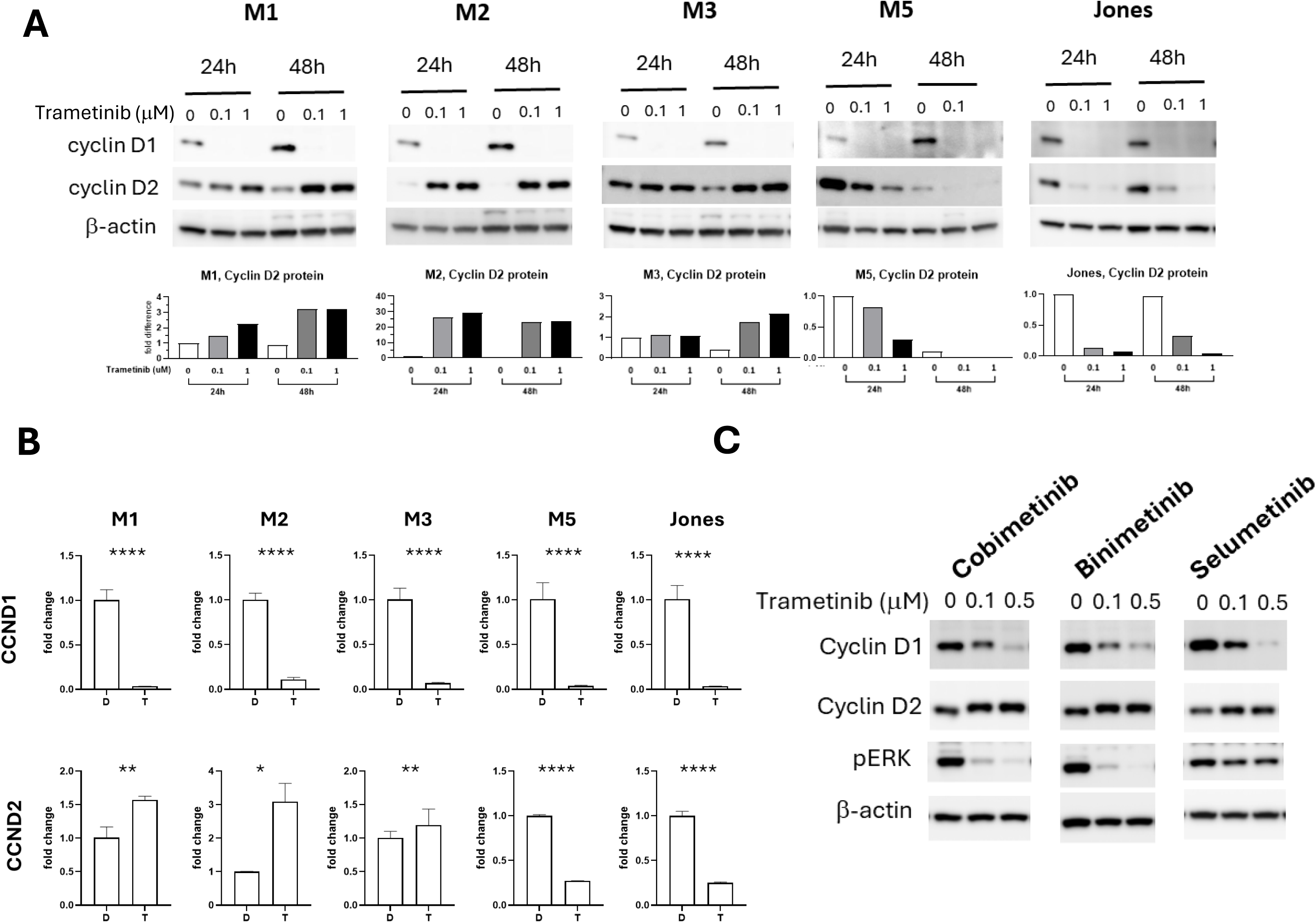
Cyclin D levels altered in canine MM cell lines upon trametinib and other MEK inhibitor exposure. A) Cyclin D1 and cyclin D2 immunoblots of canine MM cells treated with 0.1 or 1.0 μM trametinib for 24 and 48 hours. Cyclin D2 blot signal intensities were quantitated and normalized with b-actin. Expression levels are relative to no treatment control. B) CCND1 and CCND2 transcription fold change upon 1.0 μM trametinib (T) treatment for 48 hours compared to DMSO (D) treated control cultures. * p < 0.01, ** p < 0.001, **** p < 0.0001. C) Other MEK inhibitor treatments elicit similar cyclin D1 and cyclin D2 responses in M1 MM cells. M1 cells were treated with the MEK inhibitors for 48 hours at 0.1 and 0.5 μM.

The dynamic reciprocal expression of cyclin D1 and cyclin D2 in MM was further tested by knocking down (KD) cyclin D1 with siRNA. The more resistant M1 and M2 cells responded to cyclin D1 KD by increasing the expression of cyclin D2, while intrinsically more sensitive M3, M5 and Jones MM cells showed no substantive changes in cyclin D2 level when cyclin D1 expression was reduced by siRNA (Supplemental Figure S3A). In the absence of trametinib, cyclin D1 KD with siRNA by itself did not significantly alter M1 or M5 proliferative capacity (Supplemental Figure S3B), suggesting the presence of cyclin D2 was sufficient to sustain cell proliferation. Reduced proliferation was observed as a response to trametinib, although when combined with cyclin D1 KD in this series of experiments using siRNA, no further reduction in proliferation was observed (Supplemental Figure S3B).

### Inhibition of compensatory cyclin D2 expression conferred sensitivity in trametinib-resistant MM

Trametinib resistant M1 (M1T^R^) and M2 (M2T^R^) cells were generated to further investigate the role of the reciprocal expression of cyclin D1 and cyclin D2 in MM trametinib resistance. The more resistant M1T^R^ and M2T^R^ cells were adapted to trametinib by culturing the respective parental cell lines in increasing concentrations of trametinib over 2 - 4 weeks, reaching tolerance for 2 µM concentration (Figure 4A). Cyclin D1 was decreased and cyclin D2 was increased in both M1T^R^ and M2T^R^ cells, relative to the respective parental lines (Figure 4B,C,D and Supplemental Figure S4), demonstrating maintenance of the cyclin D expression pattern that was exhibited during short term drug treatment (Figure 3).

**Figure 4.**
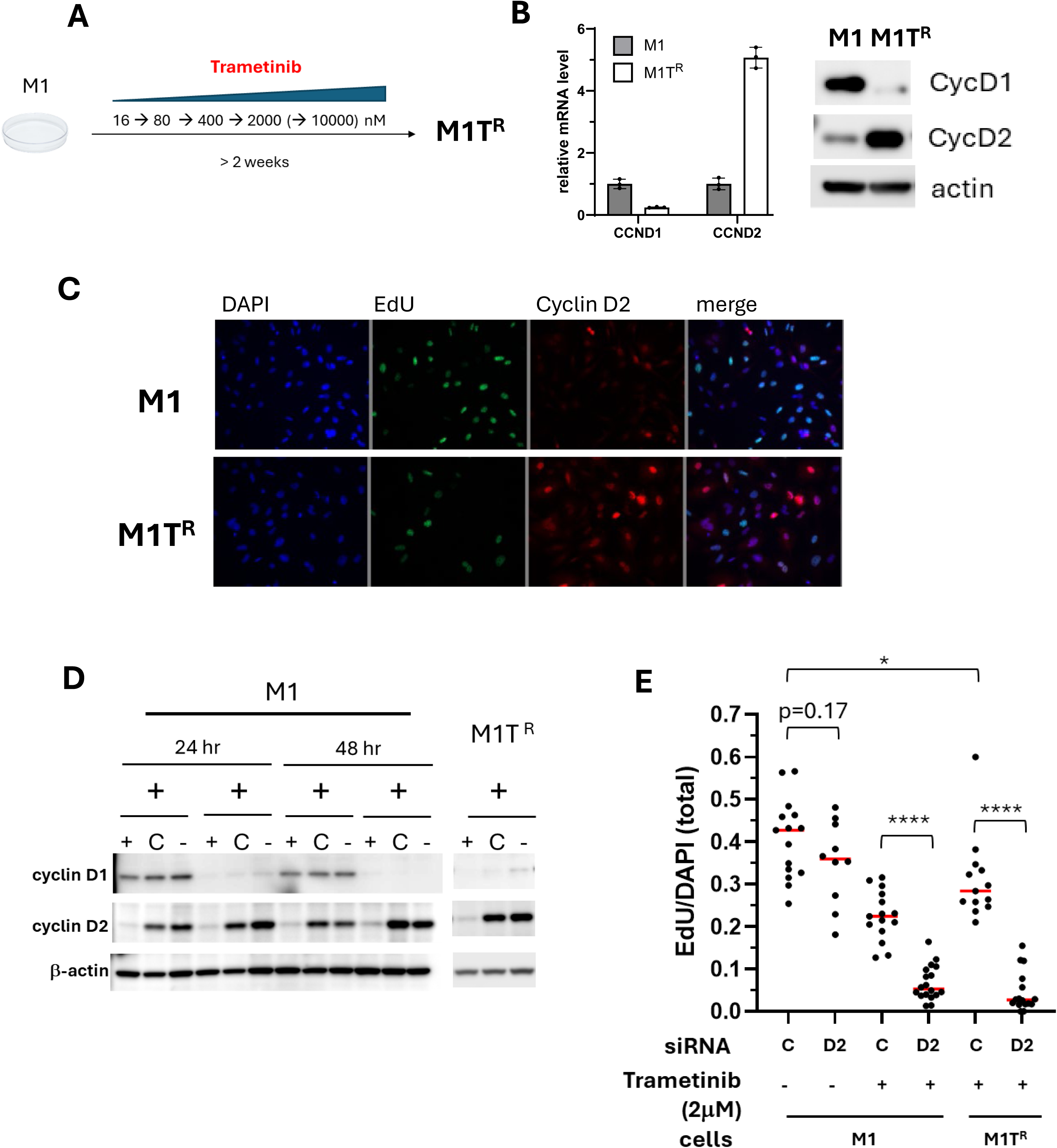
Cyclin D2 expression is critical for the proliferative capacity of trametinib-resistant M1 (M1T^R^) cells. A) M1T^R^ cells were generated by adaptive exposure of parental M1cells to increasing concentrations of trametinib for 2-4 weeks to establish the resistance phenotype. M1T^R^ cells subsequently tolerated maintenance in culture medium containing 2 μM trametinib. B) Cylin D1 was decreased in M1T^R^ cells at transcription and protein levels while cyclin D2 was increased compared to parental M1 cells. C) Immunofluorescent labeling of cyclin D2 demonstrated an increased cyclin D2 in M1T^R^ cells. The non-overlapping nuclear localization of cyclin D2 and EdU labeling corresponded to its function in cell cycle. D) Cyclin D2 was reduced in M1 and M1T^R^ cells by siRNA KD. M1 and M1T^R^ cells were transfected with CCND2 specific siRNA (D2), control siRNA (C) or no siRNA (-) for 24 hours followed by 2 μM trametinib treatment for 24 hours representing the maintenance concentration for M1T^R^ cell culture. The KD efficacy was determined by western blot. E) The impact of cyclin D2 knockdown on cell proliferation in M1 and M1T^R^ maintained in trametinib, was assessed by EdU labeling. Cell proliferation was significantly compromised by siRNA KD of cyclin D2 in trametinib-treated M1 and M1T^R^ cells. (p<0.0001).

Further examination of the potential role that compensatory cyclin D2 expression played in MM cells surviving under trametinib treatment was undertaken using siRNA to KD cyclin D2 expression. Under growth conditions lacking trametinib, cyclin D2 KD did not significantly reduce cell proliferation compared to cells transfected with control siRNA (p = 0.17), results that were similar to cyclin D1 KD alone (Figure 4E and Supplemental Figures S3 and S4B). This is consistent with our finding that canine MM cell lines appear to have the capacity to use cyclin D1 and cyclin D2 interchangeably under some circumstances (Figure 2C). Cyclin D2 KD in trametinib treated cells however, further reduced the cell proliferative capacity compared to M1 and M1T^R^ transfected with control siRNA (p < 0.0001) (Figure 4E). This outcome for M1T^R^ was similar for the adapted M2T^R^ trametinib resistant cell line (Supplemental Figure S4B). Reduced proliferative capacity in response to KD of cyclin D2 expression provided evidence that reciprocal cyclin D2 upregulation played a necessary compensatory role for maintaining cell proliferation when cyclin D1 was down regulated during trametinib exposure.

### Induced overexpression of cyclin D2 in MM confers greater trametinib resistance

To further investigate the essentiality of compensatory cyclin D2 upregulation in the observed trametinib resistance among MM cells, the trametinib-sensitive M5 cell line was engineered to express a tet-on inducible cyclin D2 (M5D2). The expressed recombinant human cyclin D2 shares 95% protein sequence homology with canine cyclin D2 (Supplemental Figure S5A) with molecular weights of 33 kD and 32.6 KD, respectively. The slightly larger recombinant cyclin D2 protein was due to the presence of a residual T2A peptide at the carboxyl-terminus (Supplemental Figure S5B, C).

Induced recombinant cyclin D2 expression in M5D2 cells was associated with diminished expression of endogenous cyclins D1 and D2 (Supplemental Figure S5B). Similar to the parental M5 cells (Figure 3A), trametinib treatment reduced the expression of endogenous cyclins D1 and D2 in M5D2 cells but had no apparent effect on induced recombinant cyclin D2 (Figure 5A). This result suggested that the decreased cyclin Ds in trametinib-treated M5 and M5D2 cells was likely due to a reduced transcription/translation of endogenous cyclins rather than increased protein degradation. Under trametinib treatment, M5D2 cells with induced exogenous cyclin D2 exhibited significantly less apoptotic cell death (Figure 5B). Resistance to cell death due to induced expression of cyclin D2 (M5D2 + doxycycline) was accompanied by superior cell proliferation (Figure 5C) and as a result, greater cell titers (Figure 5D) with more proficiently formed colonies under trametinib treatment (Figure 5E). The results indicated that induced overexpression of cyclin D2 compensated for the loss of cyclin D1 and promoted trametinib resistance in otherwise treatment-sensitive M5 MM cells that lacked an intrinsic capacity to upregulate endogenous cyclin D2 in response to trametinib (Figure 3A).

**Figure 5.**
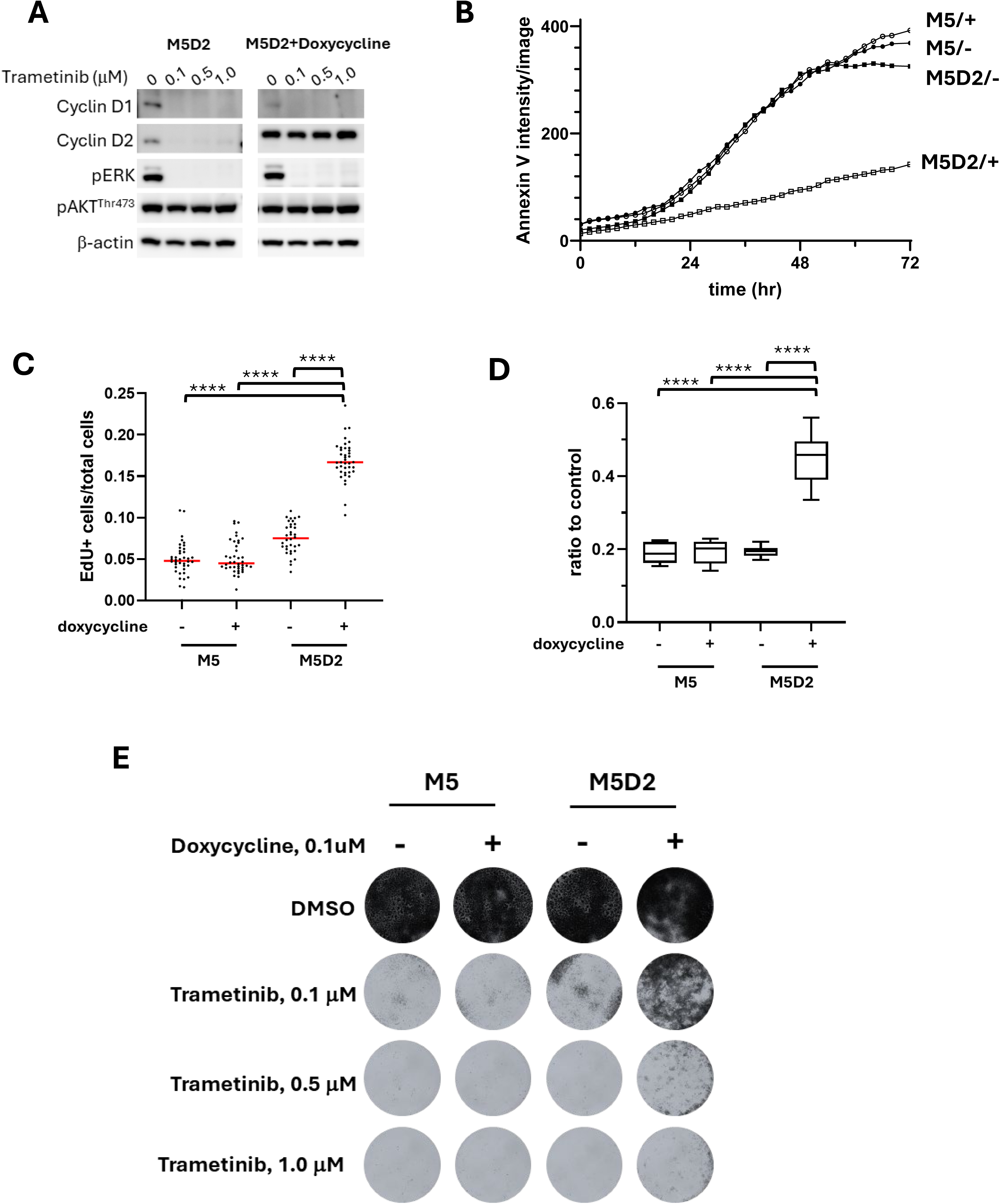
Induced cyclin D2 overexpression in M5D2 canine MM cells conferred greater resistance to the trametinib sensitive phenotype of parental M5 cells. A) The expression of cyclin D1, cyclin D2 and p-ERK in M5D2 cells with or without exogenous cyclin D2 induction. M5D2 with or without doxycycline induced exogenous cyclin D2 were treated with trametinib at 0.1, 0.5 or 1.0 μM for 48 hours. B) Annexin V-red dye was added to cells along with trametinib to monitor the cell death. The intensity of red fluorescence was measured every 2 hours for 72 hours. C) EdU incorporation was performed 48 hours after trametinib treatment. The ratios of EdU positive cells to the total cells was used to represent the proliferative activities. Each dot represents one image and each treatment was done in triplicate. **** p < 0.0001. D) MTS assay was performed to measure the cell titer 72 hours after the trametinib treatment. **** p < 0.0001. E) Colony formation assay revealed an increased cell survival in M5 cells when cyclin D2 expression was increased.

### PI3K/Akt/mTOR signal transduction pathway inhibition suppressed the trametinib-stimulated compensatory cyclin D2 expression

We previously documented that canine MMs treated with trametinib are capable of reciprocal activation of PI3K/AKT/mTOR pathway (44). In the current study, reduced ERK phosphorylation in trametinib-resistant M1T^R^ cells was accompanied by elevated AKT activation, indicating the capacity for reciprocal PI3K/Akt/mTOR signaling (Figure 6A). M1T^R^ cell reliance on PI3K/AKT pathway for survival was tested by treating M1 and M1T^R^ cells with sapanisertib, a dual mTORC1/2 inhibitor. Sapanisertib treatment reduced pAKT and pS6, inhibiting PI3K/AKT pathway signaling in both M1 and M1T^R^ cells (Figure 6B). Sapanisertib treatment led to a reduction in cyclin D1 expression (Figure 6B,C). In the case of cyclin D2, sapanisertib not only suppressed basal cyclin D2 expression in M1 cells but also inhibited the trametinib-induced upregulation of cyclin D2 in M1T^R^ and M1 cells treated with trametinib (Figure 6B,C). Notably, M1T^R^ cells were more susceptible to sapanisertib treatment than parental M1 cells (Figure 6D), further demonstrating the dependency of the conditionally adapted trametinib-resistant M1T^R^ cells on the PI3K/AKT signaling pathway for survival.

**Figure 6.**
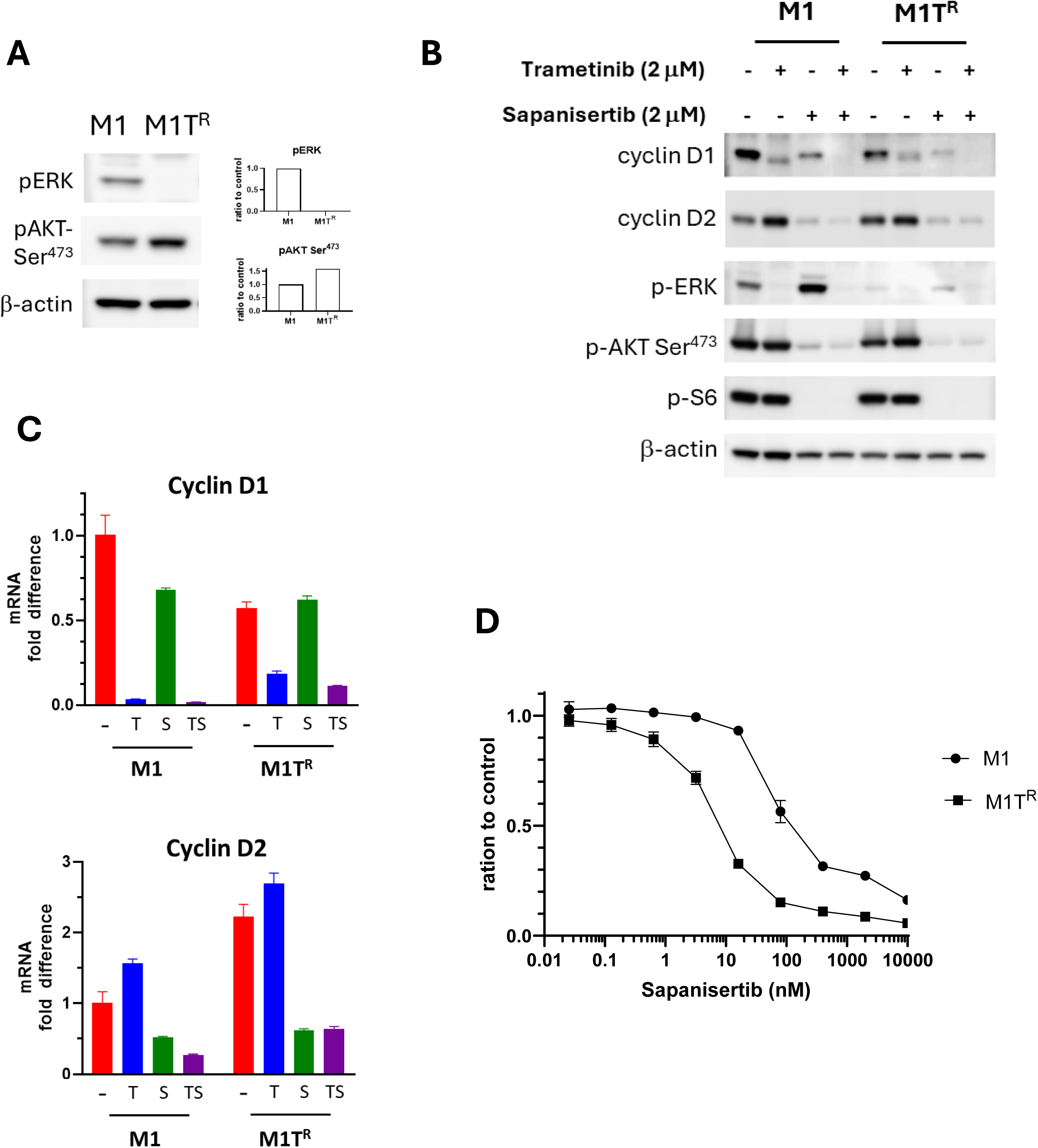
Sapanisertib, an mTORC1/2 inhibitor, countered increased cyclin D2 expression induced by trametinib, affecting survival in M1 and M1T^R^ MM. A) M1T^R^ cells with adapted trametinib resistance exhibited increased level of pAKT concurrent with pERK diminution. B) Sapanisertib alone, and in combination with trametinib, decreases levels of cyclin D1 and cyclin D2, together in association with inhibition of pAKT, and pS6 activities, the latter signaling nodes in the PI3K/AKT/mTOR pathway. C) mTORC1/2 inhibitor sapanisertib (S) inhibits cyclin D2 transcription in M1T^R^ (shown compared to treatment with trametinib (T), when combined with trametinib (TS), and to control (-)). Results are similar for M1. Sapanisertib results in less fold change reduction in CCND1 RNA transcription compared to T. D) Significantly decreased survival of M1T^R^ exposed to sapanisertib compared to parental M1 cell line (p=0.0066).

## Discussion

Despite initial treatment effectiveness, resistance to cancer therapy develops all too frequently, permitting further tumor growth and presenting a significant hurdle in achieving cancer disease remission. A variety of mutational and non-mutational events as well as changing microenvironmental factors during therapy can result in resistance mechanisms for bypassing oncogenic signaling inhibitors (35). Preservation of cancer cell proliferative capacity is a fundamental manifestation in therapeutic escape. We examined evidence for cancer cell proliferative capacity maintenance in mediating early resistance to MEK inhibition. MM cell lines with the capacity for compensatory cyclin D2 expression under trametinib treatment manifested resistance, while susceptible cells that did not upregulate cyclin D2 had significantly reduced viability. This resistance characteristic was reversed by inhibiting the activation of PI3K/AKT/mTOR pathway signaling in canine MM.

Principal oncogenic signal transduction in melanoma often engages constitutive Ras/MAPK activation and can co-involve, or recruit, dysregulated PI3K/AKT/mTOR signaling, including in the case of human and canine MM (4, 22, 45). D-type cyclins serve as important sensors for growth factors transduced by both Ras/MAPK and PI3K/AKT/mTOR pathways (46). For example, induction of cyclin D1 requires MAPK signal transduction for most cells to initiate and progress in G_1_ to S phase transitions (41). When dysregulated, overexpression or otherwise aberrant accumulation of cyclin D1 can be manifest in a variety of solid human cancers, consistent with an oncogenic function (47).

The current study results provide valuable insights into a mechanism of early resistance to trametinib linked with compensatory cyclin D expression. Through examination of five canine patient derived MM cell lines, we noted the existence of MM cell lines that were susceptible to trametinib and those that appeared relatively more intrinsically resistant. Prior to any treatment, all cell lines expressed varying levels of both cyclins D1 and D2. Trametinib reduced cyclin D1 expression across all cell lines, consistent with rational targeting expectations (48). The more trametinib sensitive cells (M5 and Jones) did not exhibit a compensatory capacity for cyclin D2 upregulation to promote proliferation, in contrast to more resistant cells (M1 and M2), which revealed upregulated cyclin D2 expression as cyclin D1 was reduced in these experiments. In fact, sensitive lines exhibited reductions of both D1 and D2 cyclins in response to trametinib. The compensatory cyclin D2 upregulation was also exclusively exhibited by the more resistant cell lines following siRNA cyclin D1 KD and appears to be a crucial and somewhat reciprocal characteristic for maintaining capacity for cell proliferation under MEK inhibitor exposure in this study.

Among the D-type cyclins, cyclin D1 and cyclin D2 generally exhibit cell-type-specific expression. Cyclin D1 is predominantly found in epithelial and certain mesenchymal cells, whereas cyclin D2 is primarily expressed in myelomonocytic/lymphoid cells and pancreatic beta cells (49, 50). This specificity suggests distinct regulatory mechanisms for each cyclin across different cell types. Yet, despite their preferential tissue distribution, cyclin D1 and cyclin D2 can functionally substitute for one another. Reciprocal expression patterns of these cyclins have been observed in various biological contexts, emphasizing their distinct yet compensatory roles in cell cycle regulation (46). For instance, cyclin D2 can compensate for the loss of cyclin D1 in estrogen-driven proliferation of uterine epithelial cells (43). In cyclin D1-null mice, cyclin D2 was able to form complexes with CDK4, translocate to the nucleus, and perform typical cyclin D1/CDK4 functions, ensuring normal cell cycle progression (43). Additionally, cyclin D2 could rescue retinal and mammary gland developmental defects in cyclin D1 knockout mice when it was knocked in to replace cyclin D1 (51). These findings indicate that although functional differences among D-type cyclins have been identified in distinct tissues, some of their functions do overlap allowing them to compensate for one another under certain conditions (24).

With evidence pointing to a capacity for reciprocal D-type cyclin substitution in MM under trametinib treatment, the fundamental function of cyclin D2 in compensating for diminished cyclin D1 in early trametinib resistance was examined. The role was confirmed through siRNA knockdown of cyclin D2 in trametinib-resistant MM cells, as well as through introduction of exogenous cyclin D2 in sensitive cells. Inhibition of cyclin D2 expression in resistant cells significantly reduced their proliferative capacity under trametinib treatment while the induced overexpression of recombinant cyclin D2 in sensitive cells promoted proliferative capacity, viability, and protection from cell death in the presence of trametinib. Accordingly, the induced exogenous cyclin D2 in trametinib-sensitive MM cells clearly performed to compensate for the reduction in endogenous cyclin D1 and cyclin D2, thereby conferring a resistance. Cyclin D2 KD by itself did not substantially affect proliferation, similar to the KD of cyclin D1 alone in the absence of trametinib.

Two-drug combinations targeting MAPK and PI3K/Akt signaling pathways in parallel impart synergistic inhibition of tumor growth and metastasis in mouse models of MM with Ras/MAPK and PI3K/AKT/mTOR activation (29, 44). This combined approach appears to be a benefit since inhibition of ERK activation by MEK inhibitors can upregulate PI3K/AKT/mTOR activation conferring resistance, as shown in the current study and previous reports (29, 44). The activation of mTOR, which is comprised of two distinct multi-protein complexes involved in regulating cellular metabolism, growth, and proliferation (mTORC1 and mTORC2), has been shown to influence regulation of cyclin D2 expression and activity (52, 53). The mTORC1 impacted cyclin D2 levels in pancreatic beta cells and T cells (52, 54). Using a controlled activation of Akt signaling and Rapamycin, an mTORC1 inhibitor, Balcazar et al. demonstrated the connection between mTORC1 activities and cyclin D2 levels in pancreatic beta cells (52). Increases in AKT activities led to increased mTORC1 activities along with elevated cyclin D2 levels, and the increased cyclin D2 was a result of enhanced protein translation and increased stability. The role of mTORC1 in cyclin D2 expression was further demonstrated in a post-partial-pancreatectomy model (55). The partial-pancreatectomy caused an increase in cyclin D2 protein levels and promoted cyclin D2 nuclear localization in an mTOR-dependent manner. Disruption of mTORC1 function in a Raptor-null mouse model revealed that mTORC1 is associated with cyclin D2 in a post-transcriptional manner and affected the stability of the cyclin D2/CDK6 complex in T cells (54). Therefore, mTORC1 regulated cyclin D2 expression primarily at the post-transcriptional level, affecting both the synthesis and stability of cyclin D2 protein.

mTORC2, on the other hand, has been shown to affect cyclin D2 expression indirectly. mTORC2 directly phosphorylates Akt at serine 473, a modification necessary for complete Akt activation. This phosphorylation event enhances Akt’s kinase activity, enabling it to phosphorylate and suppress downstream substrates, including Forkhead box O (FOXO) transcription factors (TFs) and glycogen synthase kinase 3 (GSK3) (56). FOXO TFs are known suppressors of cyclin D2 expression; the phosphorylation of FOXO TFs targets them for degradation, thereby lifting their suppressive effect on cyclin D2 transcription. GSK3β phosphorylates cyclin D2, targeting it for degradation through the ubiquitin-proteasome pathway (57). Through the activation of Akt and the subsequent inhibition of GSK3β and FOXO TFs, mTORC2 indirectly promotes the expression, stabilization, and accumulation of cyclin D2. These documented responses are consistent with finding elevated cyclin D2 expression in MM cells resistant to trametinib that could mount increased AKT activation in the face of MEK inhibition.

Congruent with these known effects of mTOR signaling, and by extension their potential to influence cyclin D2, we observed linkage between diminished cyclin D2 (and D1) level and reductions mediators of PI3K/AKT/mTOR pathway signal transduction in response to sapanisertib treatment, a dual inhibitor of mTORC1 and mTORC2. Sapanisertib’s inhibitory impact in trametinib-resistant M1T^R^ cells further supports dependency of the conditionally adapted trametinib-resistant M1T^R^ cells on the PI3K/AKT signaling pathway and cyclin D2 compensation for their survival under trametinib pressure. Therefore, inhibiting cyclin D2 may be contributing partly to the synergistic effect overcoming resistance observed, when sapanisertib was combined with MEK inhibition. Additional means for targeting potential cyclin D2 compensation may also prove beneficial.

Findings of this study substantiated that the compensatory cyclin D2 response played a role in imparting resistance by maintaining MM proliferative capacity during trametinib treatment. The results indicate the likelihood that some MM tumors have an intrinsic capacity to mount a cyclin D subtype-related compensatory response to trametinib, while other tumors would be less competent and thereby more susceptible to MEK inhibition. Current research has not sufficiently advanced to provide a molecular correlate of this characteristic and additional studies are needed. Examining the role that compensation among cyclin D subtypes plays in contributing to escape from trametinib inhibition in canine MM may provide further insight for countering resistance driven through analogous network cross talk mechanisms in this and other cancers.

## Acknowledgements

This research was supported by the Intramural Research Program, Center for Cancer Research, National Cancer Institute, Bethesda, Maryland. The authors declare that the research was conducted in the absence of any commercial or financial relationships that could be construed as a potential conflict of interest.

---

**Supplemental Figure S1.**
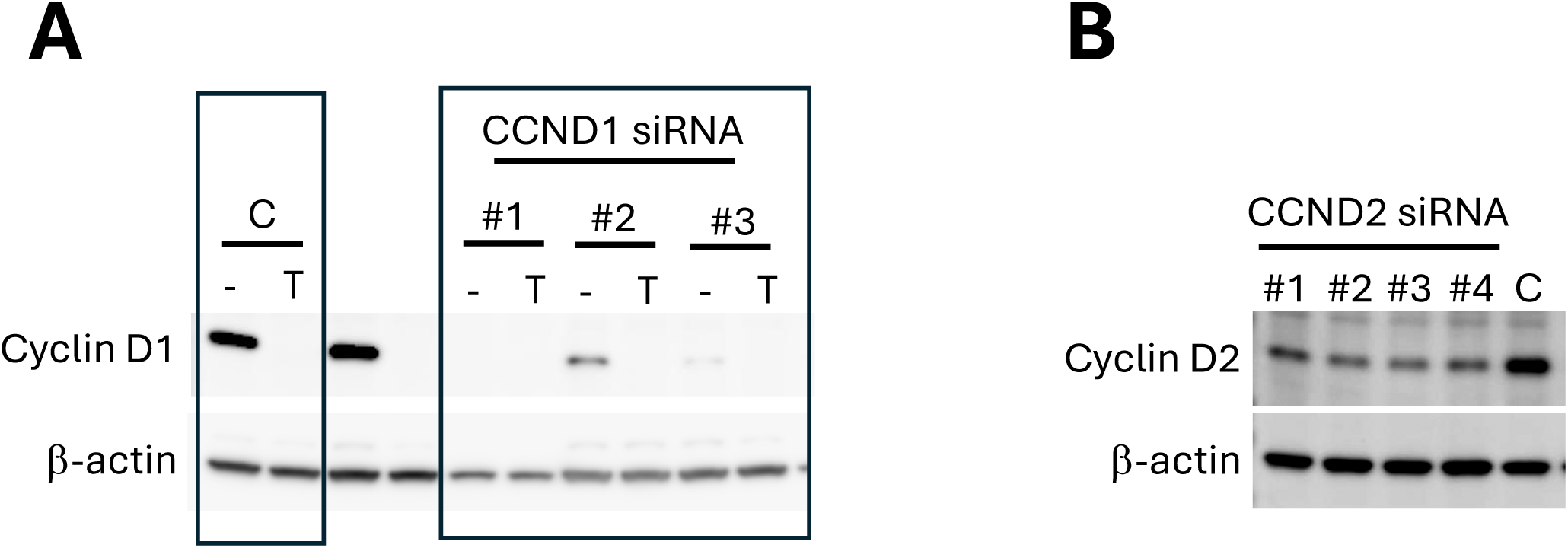
siRNA specificity test. A) Three different siRNA constructs targeting canine cyclin D1 gene (CCND1) were tested. M1 cells were transfected with 30 nM siRNA for 24 hours. The cyclin D1 level under trametinib (T) or DMSO control (-) was evaluated. B) Four different siRNA targeting cyclin D2 gene (CCND2) were similarly tested for their specificity. The cyclin D2 protein level was evaluated 24 hours after siRNA transfection.

**Supplemental Figure S2.**
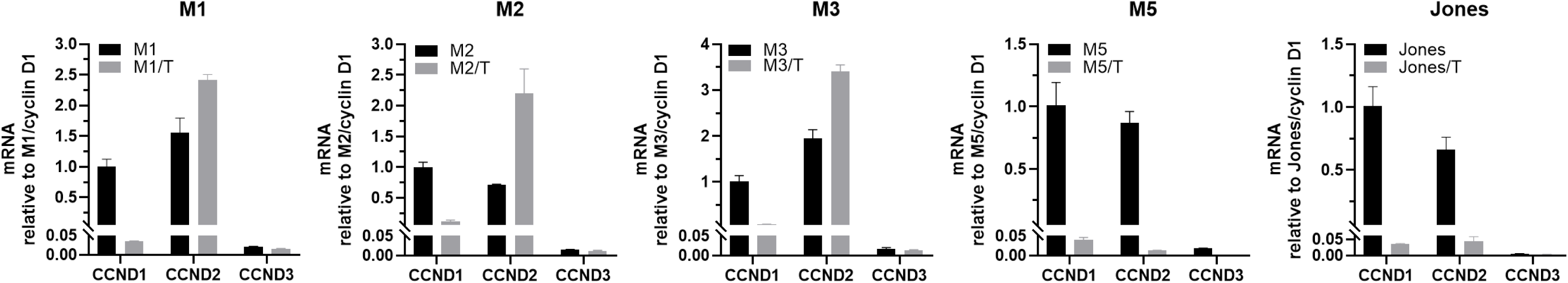
Quantitative PCR of CCND1, CCND2, and CCND3. Canine MM cells were treated with 1.0 μM trametinib (T) treatment for 48 hours. The quantity of each mRNA was reference to CCND1 level of control untreated cells in each cell line.

**Supplemental Figure S3.**
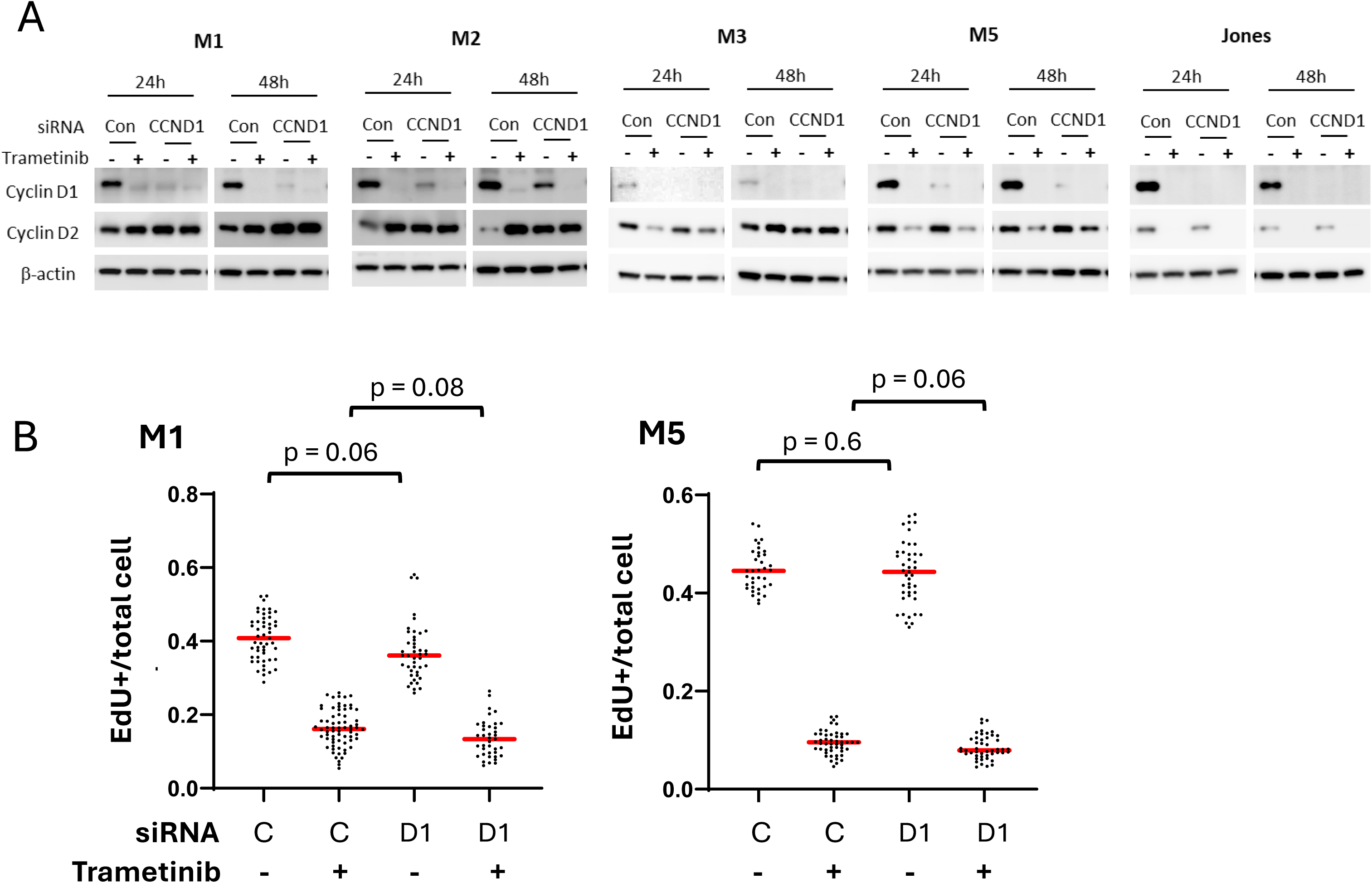
Responses of canine MM cells to cyclin D1 knock down (KD). A) Levels of cyclin D1 and D2 in canine MM cells treated with cyclin D1 siNRA followed by trametinib treatment. Cells were transfected with Cyclin D1 (CCND1) or control (Con) siRNA for 24 hours followed by 0.5 μM trametinib treatment for 24 and 48 hours. B) Cyclin D1 knock down (D1) does not significantly affect proliferative capacity in M1 and M5 cells compared to control siRNA (C), regardless of trametinib treatment. Following siRNA transfection for 24 hours, cells were treated with 0.5 μM trametinib (+) or DMSO (-) for 24 hours before EdU labeling. Each dot represents one field of view. The ratios of EdU positive cells and total cell number were used to represent the proliferative activities.

**Supplemental Figure S4.**
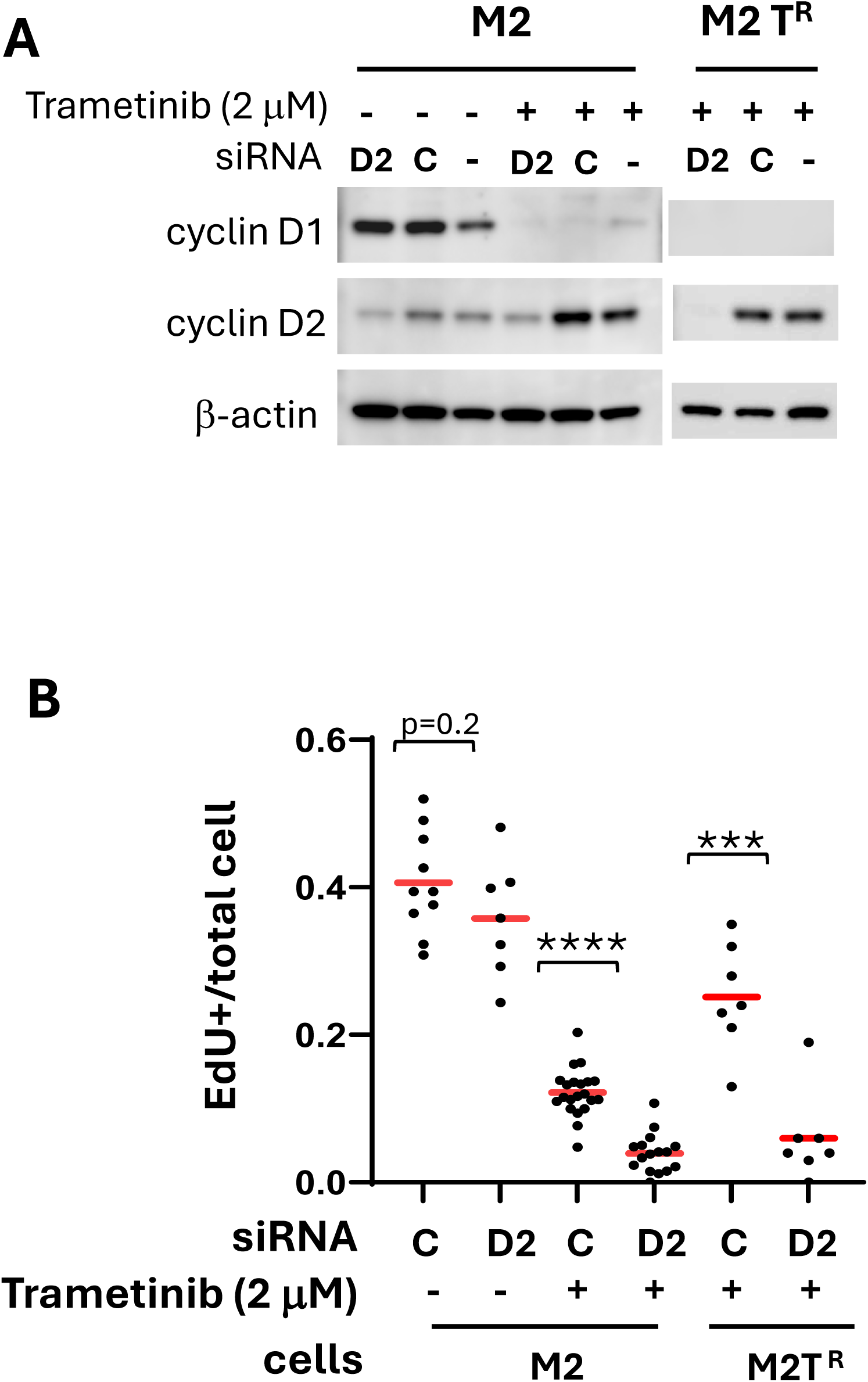
Cyclin D2 expression is similarly critical for the proliferative capacity of trametinib-resistant M2 (M2T^R^) cells. Parental M2 cells were subjected to adaptive conditioned exposure to trametinib to generate M2T^R^ trametinib resistant cells, as depicted in Figure 4A for M1T^R^. A) M2 and M2T^R^ cell were transfected with CCND2 siRNA (D2), control siRNA (C) or no siRNA (-) followed by 24 hours of 2 μM trametinib treatment (2 μM representing the maintenance concentration of trametinib for M2T^R^ cell culture). CCND2 siRNA diminished cyclin D2 level compared to expression system control (C) or no siRNA (-). B) siRNA KD of cyclin D2 in M2 and M2T^R^ cells diminished EdU incorporation indicating significantly reduced cell proliferation and increased sensitivity to trametinib.

**Supplemental Figure S5.**
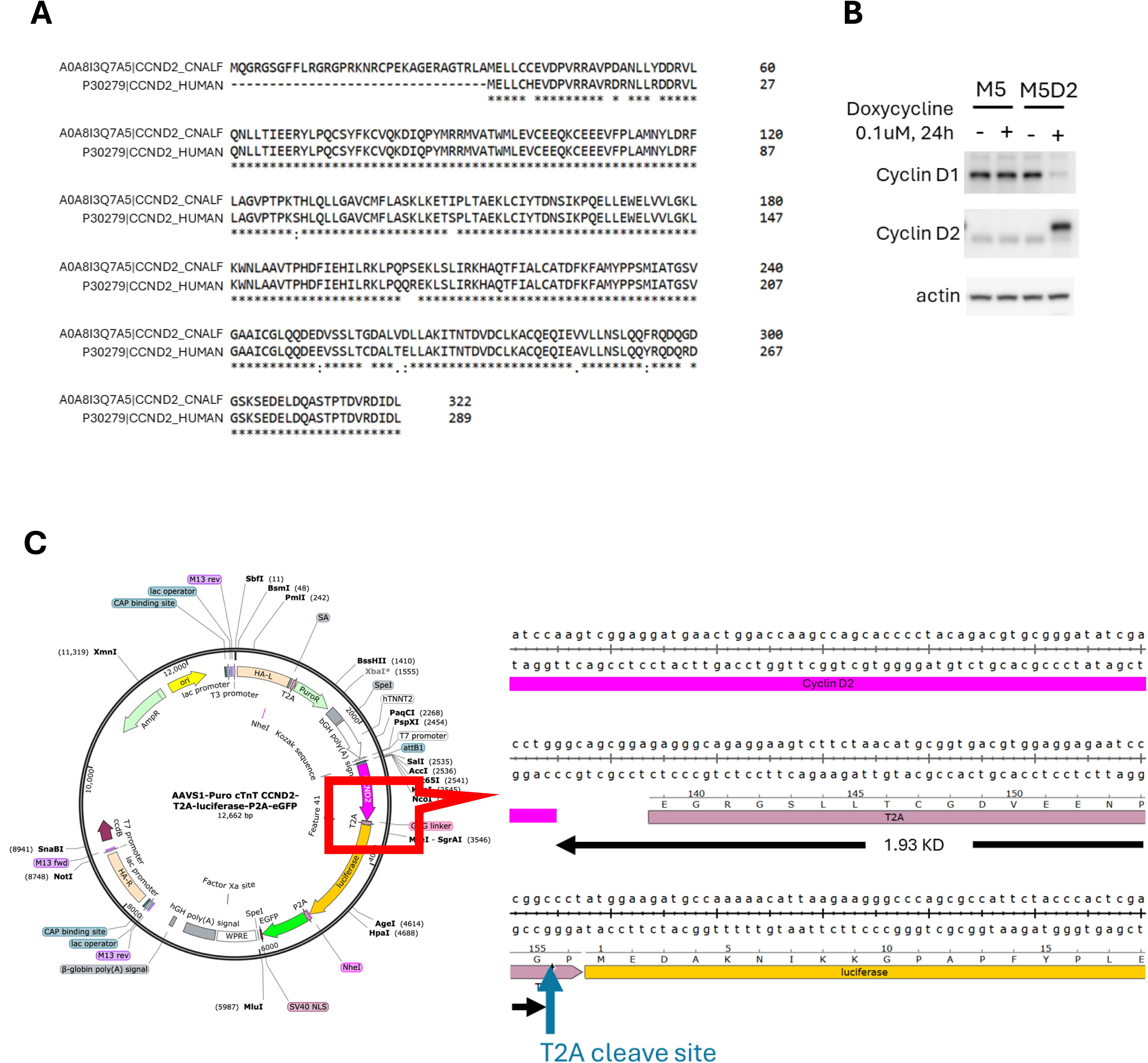
Induced stable recombinant cyclin D2 expression in canine M5 MM cells (M5D2). A) Alignment of protein sequences of human cyclin D2 (P30279) and canine cyclin D2 (A0A8I3Q7A5) revealed 95% homology. B) Western blot of cyclin D1 and D2 in M5 and M5D2 MM cells with and without the induction cyclin D2 expression. Doxycycline was added to cultures to induce the expression of recombinant cyclin D2 (+) in M5D2 cells. (C) The expression construct included a residual T2A peptide fragment at the carboxy-terminal aspect, resulting in an estimated 1.9 KD greater size molecular weight in human recombinant cyclin D2.

**Supplemental Table S1.** Reagents-siRNA, primers and antibodies siRNA.

| Target | application | vendor | catalog no. |
| --- | --- | --- | --- |
| cyclin D1 | WB, IF | cell signaling | 55506 |
| cyclin D2 | WB | cell signaling | 3741 |
| cyclin D2 | IF | Abcam | ab207604 |
| pERK | WB | cell signaling | 4270 |
| pAKT473 | WB | cell signaling | 4060 |
| actin | WB | Sigma | A5441 |

**Supplemental Table 2.** Multiple two-way comparisons between mucosal melanoma cell line viabilities over a range of trametinib concentrations.

| p values | M1 vs M2 | M1 vs M3 | M1 vs. M5 | M1 vs Jones | M2 vs M3 | M2 vs. M5 | M2 vs. Jones | M3 vs M5 | M3 vs Jones | M5 vs Jones |
| --- | --- | --- | --- | --- | --- | --- | --- | --- | --- | --- |
| 0.1 µM | 0.881 | 6.75E-04 | 1.79E-12 | 7.06E-12 | 1.75E-03 | 4.06E-08 | 1.18E-07 | 8.32E-11 | 1.09E-09 | 0.955 |
| 0.6 µM | 0.727 | 5.47E-04 | 9.72E-13 | 2.84E-13 | 3.95E-04 | 1.82E-11 | 2.30E-13 | 1.79E-05 | 2.31E-05 | 0.024 |
| 2.5 µM | 0.894 | 3.26E-06 | 7.36E-12 | 1.09E-12 | 5.14E-04 | 6.63E-06 | 5.37E-06 | 1.85E-05 | 1.61E-05 | 0.045 |
| 10.0 µM | 0.752 | 4.34E-06 | 1.26E-18 | 1.32E-17 | 8.50E-06 | 5.77E-06 | 5.14E-06 | 0.036 | 0.095 | 0.169 |

**p values correspond to data display in Figure 1A**

